# Distinct mechanisms underlying extrachromosomal telomere DNA generation in ALT cancers

**DOI:** 10.1101/2025.04.15.648952

**Authors:** Junyeop Lee, Eric J. Sohn, Jina Lee, Angelo Taglialatela, Alberto Ciccia, Jaewon Min

## Abstract

Alternative lengthening of telomeres (ALT) is a telomerase-independent telomere maintenance mechanism observed in 15% of human cancers. A hallmark of ALT cancers is the presence of C-circles, circular single-stranded DNAs (ssDNAs) enriched with cytosine-rich telomere (C-rich, CCCTAA) sequences. G-circles, containing guanosine-rich telomere (G-rich, GGGTTA) ssDNAs, also exist but are much less abundant. Recent studies indicate that excessive displacement of Okazaki fragments during lagging-strand synthesis is a unique feature of ALT telomeres and responsible for generating C-circles/C-rich ssDNAs. However, the distinct characteristics of C-circles compared to G-circles remain unclear. Here, we demonstrate that co-deficiency of the DNA translocases SMARCAL1 and FANCM leads to abundant generation of G-circle/G-rich ssDNAs. These G-rich ssDNAs mainly exist in linear form, ranging in size from 500 to 3000 nucleotides, which differs significantly from the structure and size of C-circle/C-rich ssDNAs. Mechanistically, both C-rich and G-rich ssDNAs originate from BLM/POLD-mediated excessive strand displacement; however, they differ in their origins and initiation mechanisms. Specifically, C-rich ssDNAs arise from lagging daughter strands initiated by the CST complex, whereas G-rich ssDNAs originate from leading daughter strands through RAD51-dependent G-strand synthesis. Our findings propose two distinct mechanisms for generating two different extrachromosomal telomere DNAs, C-and G-circles, during ALT-mediated telomere elongation.

## Introduction

Alternative Lengthening of Telomeres (ALT) is a telomerase-independent telomere maintenance mechanism, primarily mediated through break-induced replication (BIR) (Dilley et al. 2016; Roumelioti et al. 2016; Min et al. 2017; Sobinoff et al. 2017; Zhang et al. 2019a). ALT-positive cancers exhibit distinctive biomarkers, including extrachromosomal telomeric circular DNAs, particularly C-circles, which are circular single-stranded DNAs (ssDNAs) enriched with cytosine-rich (C-rich) sequences (CCCTAA repeats) (Cesare and Reddel 2010). C-circles serve as robust markers for ALT-positive cancers and immortalized cell lines, initially identified via phi29 polymerase-mediated rolling-circle amplification (Henson et al. 2009). Rolling-circle amplification allows for the detection of ALT cancers by magnifying C-circles in patient blood samples and tissue sections (Chen et al. 2021; Idilli et al. 2021). Currently, C-circles represent the most reliable biomarker for assessing ALT activity in human cancers. G-circles, which contain guanosine-rich (G-rich) telomere sequences (GGGTTA repeats), have also been detected using phi29 polymerase-based amplification, but are significantly less abundant compared to C-circles (Henson et al. 2009). The potential precursors of C-circles and G-circles are linear, un-circularized C-rich and G-rich ssDNAs, possibly undetectable by rolling-circle amplification. The 4SET assay, designed for the direct detection of linear C-rich ssDNAs and C-circles without phi29 polymerase-mediated amplification, is limited in detecting G-rich ssDNAs and G-circles due to their low abundance (Lee et al. 2024).

Recent studies indicate that excessive displacement of Okazaki fragments during lagging-strand synthesis is a unique characteristic of ALT telomeres (Jiang et al. 2024; Lee et al. 2024), driving the formation of C-circles and linear C-rich ssDNAs (Lee et al. 2024). However, the distinct features and detailed mechanisms underlying G-circle formation remain largely undefined due to their relatively low abundance and the absence of a suitable model system for their study.

Here, we demonstrate that co-deficiency of the DNA translocases SMARCAL1 and FANCM results in abundant production of G-circles and G-rich ssDNAs. These G-rich ssDNAs predominantly exist in a linear form and range in size from 500 to 3000 nucleotides, significantly differing from C-circles and C-rich ssDNAs. Mechanistically, although both C-rich and G-rich ssDNAs arise from BLM/POLD-mediated excessive strand displacement, they differ fundamentally in their origins and initiation mechanisms. Specifically, C-rich ssDNAs derive from lagging daughter strands initiated by the CST complex, whereas G-rich ssDNAs originate from leading daughter strands via RAD51-dependent G-strand synthesis. Our findings thus elucidate two distinct mechanisms driving the formation of extrachromosomal telomeric DNAs in ALT cancers.

## Result

### G-rich ssDNA generation under SMARCAL1 and FANCM co-deficient conditions

Mutations or loss of expression in ATRX, DAXX, and SMARCAL1 genes have been identified in ALT cancers (Heaphy et al. 2011; Lovejoy et al. 2012; Diplas et al. 2018). Loss of these genes is associated with activation of the ALT pathway and increased C-circle generation (Clynes et al. 2015; Napier et al. 2015; Diplas et al. 2018; Yost et al. 2019). Ectopic expression of ATRX protein in ATRX-deficient U2OS cells reduces the generation of C-rich ssDNA/C-circles (Clynes et al. 2015; Lovejoy et al. 2020; Jiang et al. 2024; Lee et al. 2024). To further investigate the impact of DAXX and SMARCAL1 deficiency on C-rich ssDNA generation in ALT cancers, we analyzed two ALT cancer cell lines: G292 cells, which lack wild-type (WT) DAXX expression due to a chromosomal translocation involving the DAXX gene, and NY cells, which lack SMARCAL1 expression (Fig. 1A) (Mason-Osann et al. 2018; Liu et al. 2023). We introduced a doxycycline-inducible DAXX gene expression system in G292 cells, generating G292^DAXX^ cells that express wild-type (WT) DAXX upon doxycycline treatment (Fig. S1A). Ectopic WT DAXX expression in G292 cells significantly reduced the levels of C-rich ssDNA (Fig. S1B-C), confirming that loss of DAXX expression drives ALT pathway activation and C-circle generation in G292 cells. Additionally, we established a SMARCAL1 expression system in NY cells (NY^SMARCAL1^) by expressing WT or ATPase-dead (R764Q) SMARCAL1 variants (Fig. S1D) (Ciccia et al. 2009). Expression of WT SMARCAL1 effectively reduced C-rich ssDNA generation, whereas the ATPase-dead mutant did not (Fig. S1E-F). These findings indicate that loss of SMARCAL1 expression contributes directly to C-rich ssDNA generation in NY cells, which is dependent on SMARCAL1’s ATPase activity, consistent with previous studies of SMARCAL1 function at telomeres (Poole et al. 2015; Liu et al. 2023).

**Figure 1.**
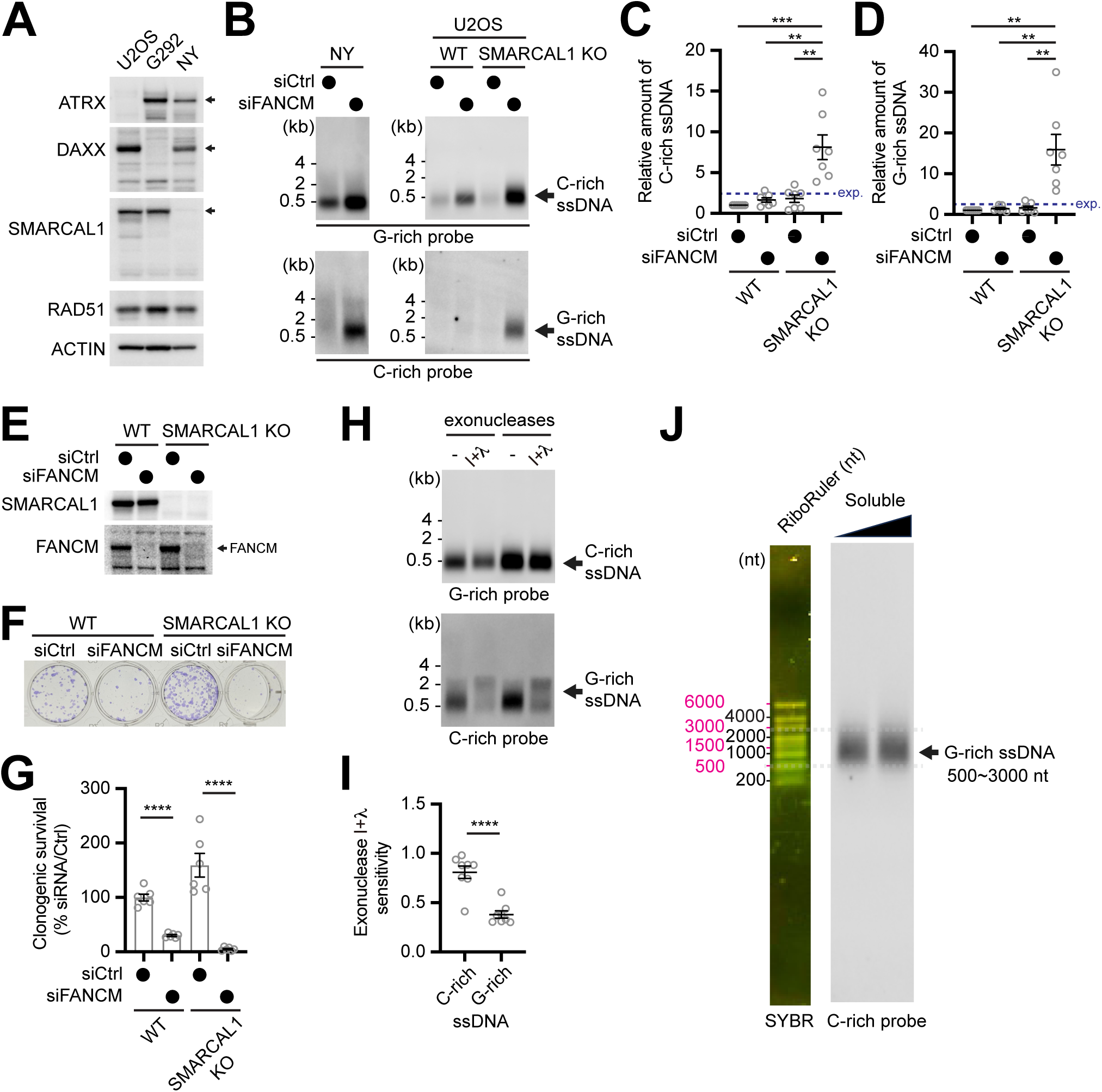
Detection of G-rich extrachromosomal telomeres. (A) Western blot analysis validating ATRX, DAXX, and SMARCAL1 protein expression levels in U2OS, G292, and NY cells. Antibodies used include anti-ATRX, anti-DAXX, anti-SMARCAL1, anti-RAD51, and anti-ACTIN (loading control). (B) Strand-specific Southern blot analysis of single-stranded extrachromosomal telomeres (4SET assay) performed on soluble DNA fractions (100 ng per sample) isolated from NY cells, U2OS wild-type (WT), and SMARCAL1 knock-out (KO) cells transfected with siRNAs targeting FANCM or control siRNAs. (C) Quantification of the assay in (B); as the relative amount of C-rich ssDNA. Blue dashed line represents the expected value for the siFANCM & SMARCAL1 KO condition. (mean ± SEM; unpaired t test). (D) Quantification of the assay in (B); as the relative amount of G-rich ssDNA. Blue dashed line represents the expected value for the siFANCM & SMARCAL1 KO condition. (mean ± SEM; unpaired t test). (E) Western blot analysis of SMARCAL1, FANCM protein expression levels in U2OS WT and SMARCAL1 KO cells after transfected with siRNAs targeting FANCM or control siRNAs. (F) Clonogenic survival of U2OS WT and SMARCAL1 KO cells after transfected with siRNAs targeting FANCM or control siRNAs. (G) Quantification of the assay in (F). (mean ± SEM; unpaired t test). (H) Nuclease assay using exonuclease I, and lambda on soluble DNA fraction of FANCM-depleted U2OS SMARCAL1 KO cells. (I) Quantification of the nuclease assay in (H); Relative amount of C-and G-rich ssDNA. (mean ± SEM; unpaired t test). (J) 4SET assay with a nucleotide ladder (RiboRuler) to determine the size of G-rich ssDNA in FANCM-depleted U2OS SMARCAL1 KO cells.

Next, we investigated the effect of depleting FANCM, a known suppressor of the ALT pathway whose loss results in significant C-circle and C-rich ssDNA generation. Multiple studies have demonstrated that FANCM depletion in U2OS cells leads to increased levels of C-rich ssDNA and C-circles (Lu et al. 2019; Pan et al. 2019; Silva et al. 2019; Lee et al. 2024). To extend this observation, we depleted FANCM using siRNA in NY cells lacking SMARCAL1 expression. FANCM depletion in NY cells not only increased C-rich ssDNA generation (Fig. 1B; top left) but also unexpectedly resulted in the appearance of G-rich ssDNA (Fig. 1B; bottom left). We hypothesized that G-rich ssDNA formation upon FANCM depletion in NY cells resulted from co-deficiency with SMARCAL1. To test this hypothesis, we compared G-rich ssDNA abundance in WT U2OS cells and SMARCAL1 knockout (KO) U2OS cells after FANCM depletion. U2OS SMARCAL1 KO clones were generated with low efficiency due to the dependency of ALT cells on the SMARCAL1 gene (Taglialatela et al., submitted) (Lu and Pickett 2022); however, a few clones adapted to proliferate, enabling us to successfully obtain SMARCAL1 KO clones. Consistent with previous findings (Lu et al. 2019; Lee et al. 2024), FANCM depletion in WT U2OS cells increased only C-rich ssDNA generation (Fig. 1B; G-rich probe) whereas G-rich ssDNA remained undetectable (Fig. 1B; C-rich probe). In contrast, FANCM depletion in SMARCAL1 KO U2OS cells led to substantial increases in both C-rich and G-rich ssDNA (Fig. 1B; bottom right, 1C-D). Depletion efficiencies of FANCM and SMARCAL1 expression levels were confirmed by western blot (Fig. 1E). Both WT and SMARCAL1 KO U2OS cells exhibited reduced proliferation upon FANCM depletion (Fig. 1F-G), indicating that the generation of G-rich ssDNA was not simply due to impaired proliferation.

To further confirm these findings, we introduced a doxycycline-inducible WT SMARCAL1 expression system in SMARCAL1 KO U2OS cells (Fig. S1G). SMARCAL1 re-expression significantly reduced both C-rich ssDNA and prevented G-rich ssDNA formation upon FANCM depletion (Fig. S1H-J). Similar results were observed when expressing WT SMARCAL1 in NY^SMARCAL1^ cells (Fig. S1K-M). We also validated FANCM depletion using CRISPR/Cas9-mediated genome editing with gRNAs targeting FANCM (Fig. S1N), observing increased G-rich ssDNA in SMARCAL1 KO cells (Fig. S1O) and proliferation defects in both WT and SMARCAL1 KO U2OS cells (Fig. S1P-Q). Collectively, these data demonstrate that co-deficiency of SMARCAL1 and FANCM promotes the generation of G-rich ssDNA in ALT cells.

We further investigated the mechanism of FANCM in suppressing G-rich ssDNA generation. FAAP24, a FANCM-interacting protein, recruits FANCM to DNA intermediate structures formed during DNA replication and repair (Ciccia et al. 2007). FAAP24 depletion is known to increase C-circle levels similarly to FANCM depletion (Pan et al. 2019). We examined whether FAAP24 depletion under SMARCAL1 deficiency conditions also leads to G-rich ssDNA generation. Indeed, FAAP24 depletion increased both C-rich and G-rich ssDNA in NY cells (Fig. S1R) and U2OS SMARCAL1 KO cells (Fig. S1S), suggesting that FANCM’s function in association with FAAP24 is crucial for preventing G-rich ssDNA generation.

Lastly, we characterized the structures of C-rich and G-rich ssDNAs generated under SMARCAL1 and FANCM deficiency conditions using exonuclease assays. G-rich ssDNA was highly sensitive to exonuclease I and lambda exonuclease treatment (Fig. 1H, bottom), whereas C-rich ssDNA exhibited greater resistance (Fig. 1H, top). Overall, G-rich ssDNA was more sensitive to exonuclease digestion compared to C-rich ssDNA (Fig. 1I), indicating that G-rich ssDNA predominantly exists in a linear form rather than circular. Size analysis using a nucleotide ladder indicated G-rich ssDNAs ranged from 500 to 3000 nucleotides (Fig. 1J). This is longer than C-rich ssDNAs, which are from 200 to 1500 nucleotides (Lee et al. 2024). These results indicate that G-rich ssDNAs exhibit unique features distinct from those of C-rich ssDNAs.

### Telomere clustering and RPA accumulation under SMARCAL1 and FANCM co-deficient conditions

Telomere clustering is a hallmark of the ALT pathway (Yeager et al. 1999; Cesare and Reddel 2010; Heaphy et al. 2011; Min et al. 2019; Zhang et al. 2020; Zhang et al. 2021). Previous studies show that depletion of SMARCAL1 or FANCM in ALT cancers results in telomere clustering, visualized as large telomere foci (Cox et al. 2016; Pan et al. 2017; Silva et al. 2019). To determine whether the presence of these large telomere foci (>0.4 micrometer) indicates ongoing telomere clustering, we compared cells with and without large telomere foci (Fig. S2A). Telomere clustering results in distances between foci that differ significantly from random distributions, which can be quantitatively assessed using spatial analysis plugins in the Icy software, specifically Ripley’s K-function and nearest neighbor analysis (Fig. S2B). We excluded large foci from the analysis when these large foci could be counted as multi-foci, potentially interfering with our analysis. Our analysis of telomere foci in cells containing large telomere foci revealed clustering above the defined threshold, distinct from cells without large telomere foci (Fig. S2C). These results indicate that cells exhibiting large telomere foci reliably represent ongoing telomere clustering events in ALT cancers.

Based on this methodology, we analyzed telomere foci in FANCM-depleted U2OS WT and SMARCAL1 KO cells. Upon FANCM depletion, U2OS SMARCAL1 KO cells exhibited a significant increase in the number of large telomere foci per cell (Fig. 2A-B), with over 40% of cells showing large telomere foci (Fig. S2D).

**Figure 2.**
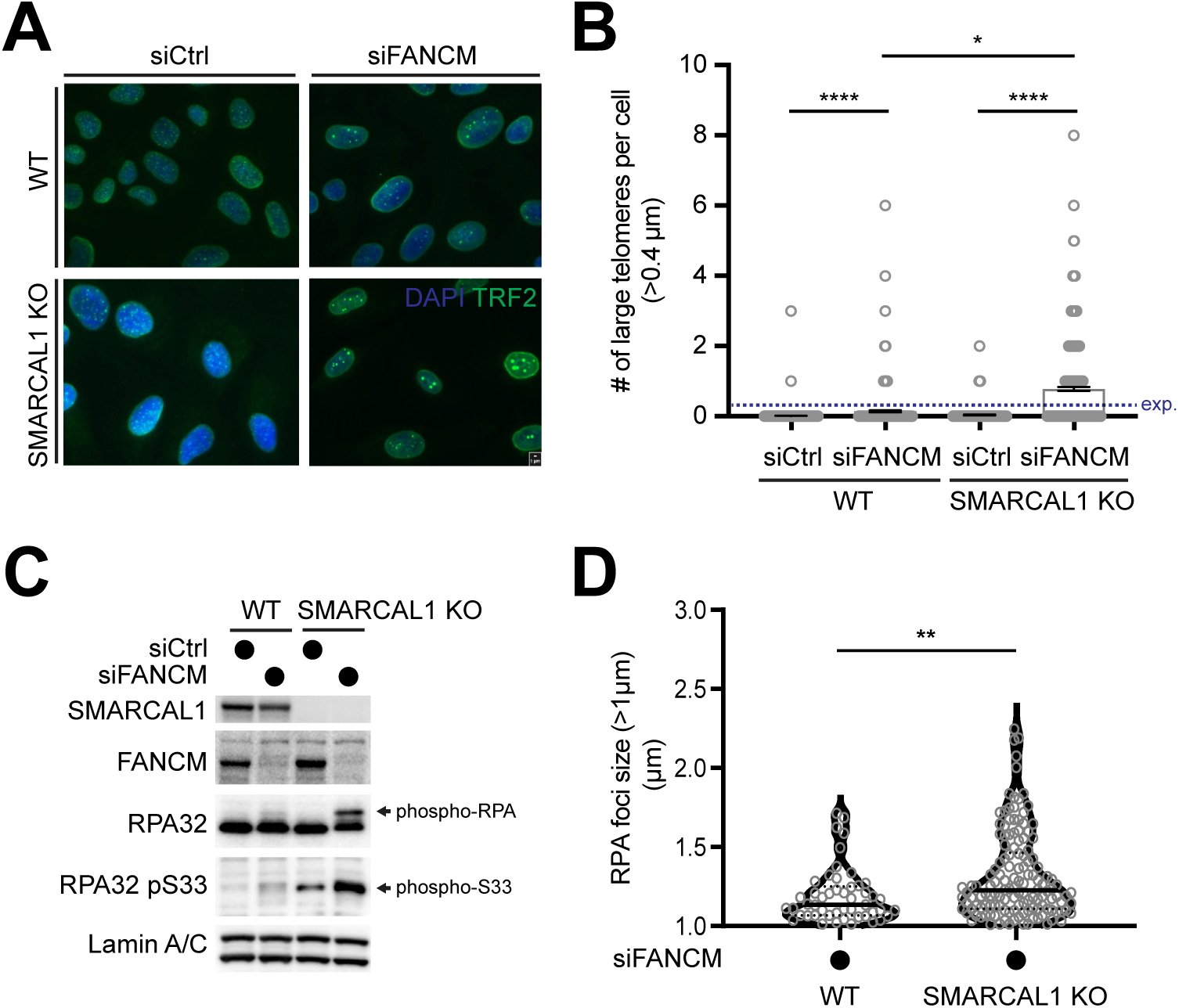
Telomere clustering and RPA phosphorylation in FANCM and SMARCAL1 co-deficient U2OS cells. (A) Immunofluorescence staining of TRF2 (green) and DAPI (blue) in U2OS WT and SMARCAL1 KO cells after transfected with siRNAs targeting FANCM or control siRNAs. (B) Quantification of the number of large telomeres (more than 0.4 micrometer) per cells. Blue dashed line represents the expected value for the siFANCM & SMARCAL1 KO condition. (mean ± SEM; unpaired t test). (C) Western blot analysis of SMARCAL1, FANCM, RPA32, phosphorylated Serine 33 in RPA32 (RPA32 pS33), Lamin A/C (loading control) in U2OS WT and SMARCAL1 KO cells after transfected with siRNAs targeting FANCM or control siRNAs. (D) Quantification of the large RPA foci size (more than 1 micrometer) in FANCM depleted U2OS WT and SMARCAL1 KO cells. (median, Mann-Whitney test)

Additionally, phosphorylated RPA was observed in FANCM-depleted SMARCAL1 KO cells, with RPA32 phosphorylation detected specifically at serine 33 (Fig. 2C). Phosphorylation of RPA32 at S33 increased synergistically following co-depletion of FANCM and SMARCAL1 (Fig. S2E). RPA phosphorylation potentially indicates gap formation resulting from FANCM and SMARCAL depletion (Vassin et al. 2009). FANCM depletion resulted in the formation of large RPA foci (Fig. S2F). Both the number of large RPA foci (Fig. S2G) and the percentage of cells containing these large RPA foci (Fig. S2H) increased in FANCM and SMARCAL1 co-depleted cells. Notably, the size of individual RPA foci also increased upon co-depletion of FANCM and SMARCAL1 (Fig. 2D). Co-depletion of FANCM and SMARCAL1 leads to gap formation accompanied by telomere clustering.

### BLM/POLD-mediated strand displacement generates G-rich ssDNAs, same as C-rich ssDNAs

Next, we investigated the underlying mechanisms responsible for G-rich ssDNA generation. First, we examined whether excessive strand displacement mediated by BLM and POLD is essential for G-rich ssDNA production, as previously established for C-rich ssDNAs (Dilley et al. 2016; Sobinoff et al. 2017; Min et al. 2019; Loe et al. 2020; Jiang et al. 2024; Lee et al. 2024). Depletion of POLD3, a subunit of POLD, in FANCM-depleted U2OS SMARCAL1 KO cells significantly reduced the levels of both C-rich (Fig. 3A-B) and G-rich ssDNAs (Fig. 3A, C). Similarly, BLM depletion in U2OS SMARCAL1 KO cells also decreased the levels of both C-rich (Fig. 3D-E) and G-rich ssDNAs (Fig. 3D, F).

**Figure 3.**
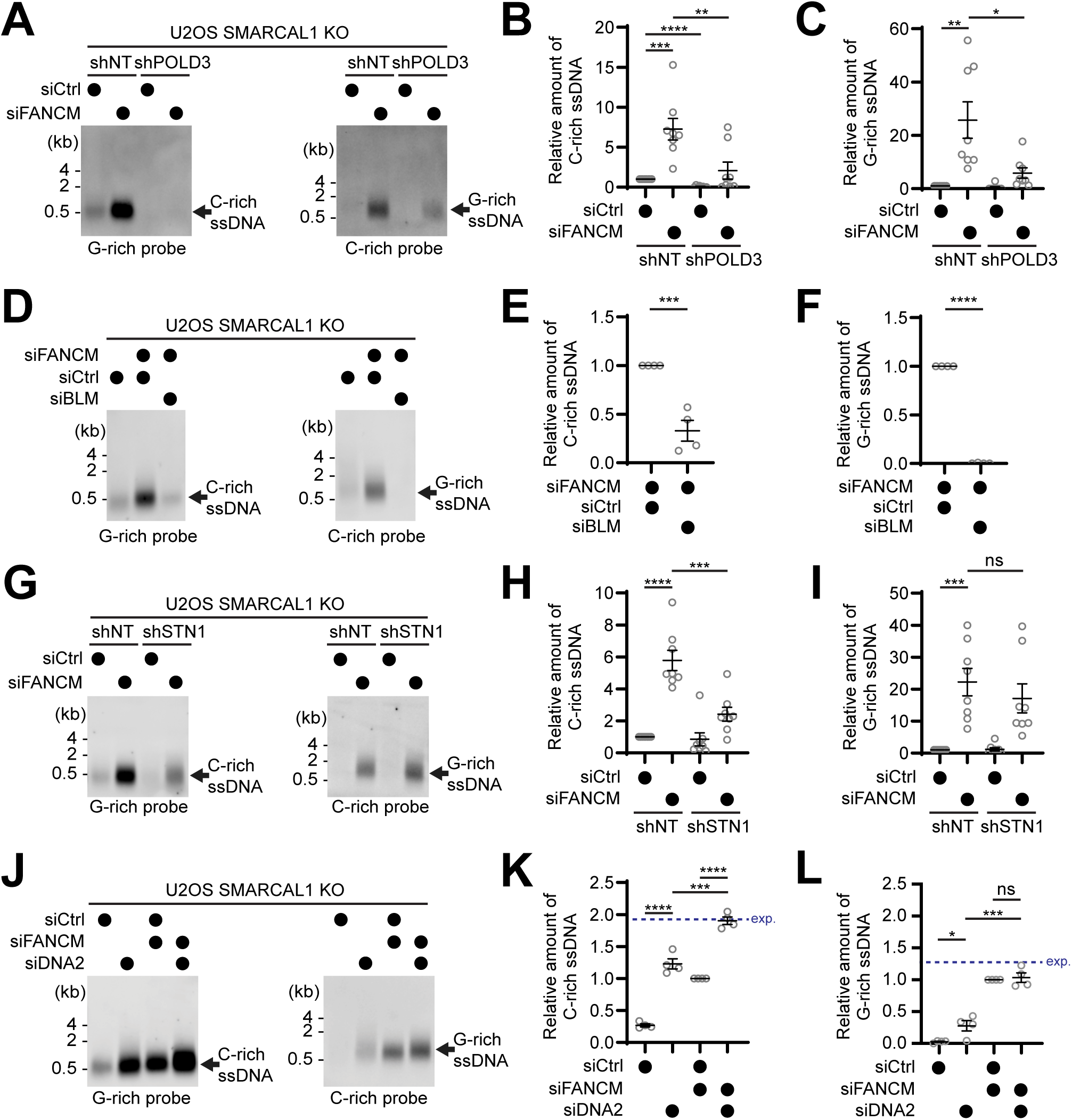
Distinct mechanisms of C-rich and G-rich ssDNA generation. (A) 4SET assay with POLD3-depleted (shPOLD3) and control (shNT) infected U2OS SMARCAL1 KO cells after transfected with siRNAs targeting FANCM or control siRNAs. (B) Quantification of the assay in (A); as the relative amount of C-rich ssDNA. (mean ± SEM; unpaired t test). (C) Quantification of the assay in (A); as the relative amount of G-rich ssDNA. (mean ± SEM; unpaired t test). (D) 4SET assay with U2OS SMARCAL1 KO cells after transfected with siRNAs targeting FANCM, BLM, and control siRNAs. (E) Quantification of the assay in (D); as the relative amount of C-rich ssDNA. (mean ± SEM; unpaired t test). (F) Quantification of the assay in (D); as the relative amount of G-rich ssDNA. (mean ± SEM; unpaired t test). (G) 4SET assay with STN1-depleted (shSTN1) and control (shNT) infected U2OS SMARCAL1 KO cells after transfected with siRNAs targeting FANCM or control siRNAs. (H) Quantification of the assay in (G); as the relative amount of C-rich ssDNA. (mean ± SEM; unpaired t test). (I) Quantification of the assay in (G); as the relative amount of G-rich ssDNA. (mean ± SEM; unpaired t test). (J) 4SET assay with U2OS SMARCAL1 KO cells after transfected with siRNAs targeting FANCM, DNA2, and control siRNAs. (K) Quantification of the assay in (J); as the relative amount of C-rich ssDNA. Blue dashed line represents the expected value for the siFANCM & siDNA2 condition. (mean ± SEM; unpaired t test). (L) Quantification of the assay in (J); as the relative amount of G-rich ssDNA. Blue dashed line represents the expected value for the siFANCM & siDNA2 condition. (mean ± SEM; unpaired t test).

We validated these findings in NY cells. Initially, we confirmed that C-rich ssDNA generation in NY cells is dependent on POLD and BLM helicase activities. Depletion of POLD3 in NY cells using shRNA targeting POLD3 (Fig. S3A) led to decreased C-rich ssDNA generation (Fig. S3B-C). Similarly, depletion of BLM in NY cells reduced the generation of C-rich ssDNAs (Fig. S3D-F). Furthermore, BLM depletion in FANCM-depleted NY cells also reduced both C-rich (Fig. S3G-H) and G-rich ssDNAs (Fig. S3G, S3I). These results collectively indicate that G-rich ssDNAs, in the same manner as C-rich ssDNAs, originate from strand displacement processes.

SLX4 antagonizes BLM-mediated dissolution by resolving D-loops at telomeres (Sarkar et al. 2015). Since SLX4 depletion increases C-circles in ALT cells (Sobinoff et al. 2017), we tested whether SLX4 depletion also elevates the generation of C-rich and G-rich ssDNAs. We depleted SLX4 using siRNA (Fig. S3J). SLX4 depletion increased levels of both C-rich (Fig. S3K-L) and G-rich ssDNAs (Fig. S3K, S3M), confirming that G-rich ssDNA generation is BLM-dependent strand displacement.

### The CST complex and DNA2 flap nuclease are dispensable for G-rich ssDNA generation, unlike their roles in C-rich ssDNA generation

We next tested whether the CST complex, essential for generating C-rich ssDNAs by initiating additional Okazaki fragments (Huang et al. 2017; Lee et al. 2024), also contributes to G-rich ssDNA production. By introducing shRNA targeting STN1, a component of the CST complex, we found that STN1 depletion significantly reduced C-rich ssDNA levels (Fig. 3J-K) but did not affect G-rich ssDNA levels (Fig. 3J, L). We confirmed these findings using siRNA targeting STN1 (Fig. S3N). Transient depletion of STN1 with siRNA similarly decreased C-rich ssDNA (Fig. S3O-Q), but unexpectedly increased G-rich ssDNA generation in FANCM-depleted U2OS SMARCAL1 KO cells. These data indicate that G-rich ssDNA generation is independent of the CST complex.

Finally, we investigated DNA2, known to suppress C-rich ssDNA generation by processing flap structures of Okazaki fragments (O’Sullivan et al. 2014; Jiang et al. 2024; Lee et al. 2024). Co-depletion of DNA2 and FANCM in U2OS SMARCAL1 KO cells additively increased C-rich ssDNA levels (Fig. 3J-K) but did not further enhance G-rich ssDNA generation (Fig. 3J, 3L), demonstrating that DNA2 does not process G-rich ssDNAs.

Collectively, our data indicate that the CST complex is required for generating C-rich ssDNAs but is dispensable for G-rich ssDNA synthesis. Both C-rich and G-rich ssDNA generation rely on POLD/BLM-mediated strand displacement, while DNA2-mediated 5’ flap processing exclusively influences C-rich ssDNA formation. Given that CST and DNA2 primarily associate with lagging strand, we reason that C-circles originate from lagging daughter strands, whereas G-circles arise from leading daughter strands. However, the initiation mechanism for G-rich ssDNA synthesis appears distinct, given its independence from the CST complex. Therefore, we next sought to determine the initiation mechanisms underlying G-rich ssDNA generation.

### RAD51 dependent G-rich ssDNA generations

RAD51 is involved in HR (homologous recombination)-dependent gap-filling processes during the S phase (Adar et al. 2009; Piberger et al. 2020; Tirman et al. 2021). We investigated whether RAD51 is required for generating G-rich ssDNA by depleting RAD51 and PARI (a human homolog of yeast Srs2, anti-recombinase) using siRNAs (Moldovan et al. 2012). RAD51 depletion in FANCM-depleted U2OS SMARCAL1 KO cells did not alter C-rich ssDNA generation (Fig. 4A-B; lane 2). However, RAD51 depletion in FANCM-depleted U2OS cells increased C-rich ssDNA levels (Fig. S4A-B). In contrast, PARI depletion decreased C-rich ssDNA in both FANCM-depleted and undepleted U2OS SMARCAL1 KO cells (Fig. 4A-B; lanes 3, 6). Strikingly, RAD51 depletion in FANCM-depleted U2OS SMARCAL1 KO cells significantly reduced G-rich ssDNA by around 50% (Fig. 4A, C; lane 5), whereas PARI depletion significantly increased G-rich ssDNA levels (Fig. 4A, C; lane 6). These results suggest RAD51 is required for the initiation of G-rich ssDNA formation.

**Figure 4.**
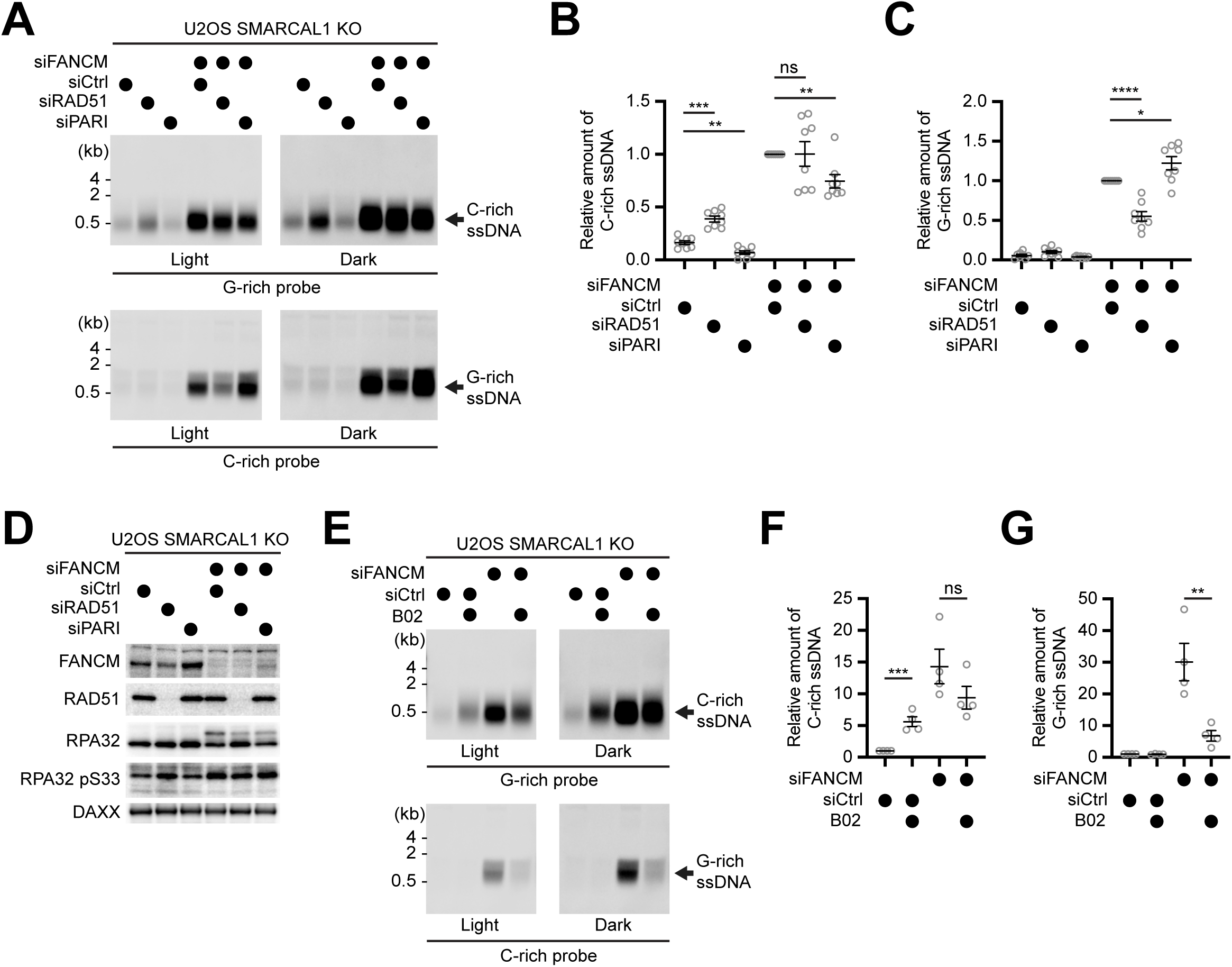
RAD51-dependent G-rich ssDNA generation. (A) 4SET assay with U2OS SMARCAL1 KO cells after transfected with siRNAs targeting FANCM, RAD51, PARI, and control siRNAs. (B) Quantification of the assay in (A); as the relative amount of C-rich ssDNA. (mean ± SEM; unpaired t test). (C) Quantification of the assay in (A); as the relative amount of G-rich ssDNA. (mean ± SEM; unpaired t test). (D) Western blot analysis of RAD51, FANCM, RPA32, phosphorylated Serine 33 in RPA32 (RPA32 pS33), DAXX (loading control) in U2OS SMARCAL1 KO cells after transfected with siRNAs targeting FANCM, RAD51, PARI, and control siRNAs. (E) 4SET assay with U2OS SMARCAL1 KO cells treated with or without B02 (RAD51 inhibitor) after transfected with siRNAs targeting FANCM, or control siRNAs. (F) Quantification of the assay in (E); as the relative amount of C-rich ssDNA. (mean ± SEM; unpaired t test). (G) Quantification of the assay in (E); as the relative amount of G-rich ssDNA. (mean ± SEM; unpaired t test).

We also evaluated RPA phosphorylation and the expression levels of FANCM, RAD51, and PARI (Fig. 4D, S4C-D). PARI depletion in U2OS SMARCAL1 KO cells increased RAD51 foci formation (Fig. S4E), consistent with previous observation (Moldovan et al. 2012). RAD51 depletion in U2OS SMARCAL1 KO cells increased RPA32 S33 phosphorylation (Fig. 4D, S4C; lane 2), potentially indicating increased replication gap formation, consistent with RAD51’s role in suppressing replication gaps (Hashimoto et al. 2010; Kolinjivadi et al. 2017; Mann et al. 2022). We hypothesized that increased C-rich ssDNA in RAD51-depleted U2OS SMARCAL1 KO cells could be due to CST-mediated C-strand synthesis on lagging strands. Indeed, STN1 depletion using shRNA reduced the elevated C-rich ssDNA levels observed upon RAD51 depletion (Fig. S4F-G; lanes 2, 5). This result indicates that RAD51 loss generates gaps on lagging strands, which triggers CST-dependent C-strand synthesis and thus increases C-rich ssDNA generation.

Moreover, PARI depletion resulted in reduced C-rich ssDNA generation, similar to STN1 depletion (Fig. S4F-G; lanes 3-4). Importantly, co-depletion of PARI and STN1 did not further reduce C-rich ssDNA levels (Fig. S4F-G; lane 6), confirming that the increased C-rich ssDNA generation caused by RAD51 depletion depends on the CST complex.

Next, we examined RAD51-dependent G-rich ssDNA generation using B02, a small-molecule inhibitor of RAD51 (Huang et al. 2011). Treatment with B02 resulted in a dramatic reduction of G-rich ssDNA in FANCM-depleted SMARCAL1 KO cells (Fig. 4E, G; lanes 3-4), while C-rich ssDNA levels remained largely unchanged (Fig. 4E-F; lanes 3-4). These results confirm that G-rich ssDNA generation under FANCM and SMARCAL1 co-deficient conditions depends on RAD51 recombinase activity. Interestingly, B02 treatment alone increased C-rich ssDNA generation (Fig. 4E-F; lanes 1-2), consistent with our observations in RAD51-depleted samples using siRNA (Fig. 4A-B; lanes 1-2). Similarly, RAD51 inhibition in NY cells significantly elevated C-rich ssDNA levels by five-fold (Fig. S4H-I). This increase in C-rich ssDNA upon RAD51 inhibition was mitigated by STN1 depletion (Fig. S4J-K), confirming that CST-dependent C-strand synthesis is responsible for the observed elevation in C-rich ssDNA under RAD51 inhibition conditions.

### Generation of G-rich ssDNA can be suppressed by anti-recombinases

SMARCAL1, ZRANB3, and HLTF are related members of the SNF2 family, which play specialized roles in maintaining genome stability during DNA replication, particularly by promoting fork reversal in response to replication stress (Adolph and Cortez 2024). We investigated whether the generation of G-rich ssDNA observed upon FANCM deficiency is specific to SMARCAL1 or if deficiencies in ZRANB3 and HLTF could produce similar effects. We introduced gRNAs targeting HLTF or ZRANB3 to induce their depletion (Fig. 5A). Depletion of HLTF or ZRANB3 alone did not significantly generate G-rich ssDNA compared to SMARCAL1 KO cells (Fig. S5A-C). Although ZRANB3 depletion led to a modest (∼2-fold) increase, this was minimal compared to the substantial (∼11-fold) increase observed in SMARCAL1 KO cells (Fig. S5C, 1D).

**Figure 5.**
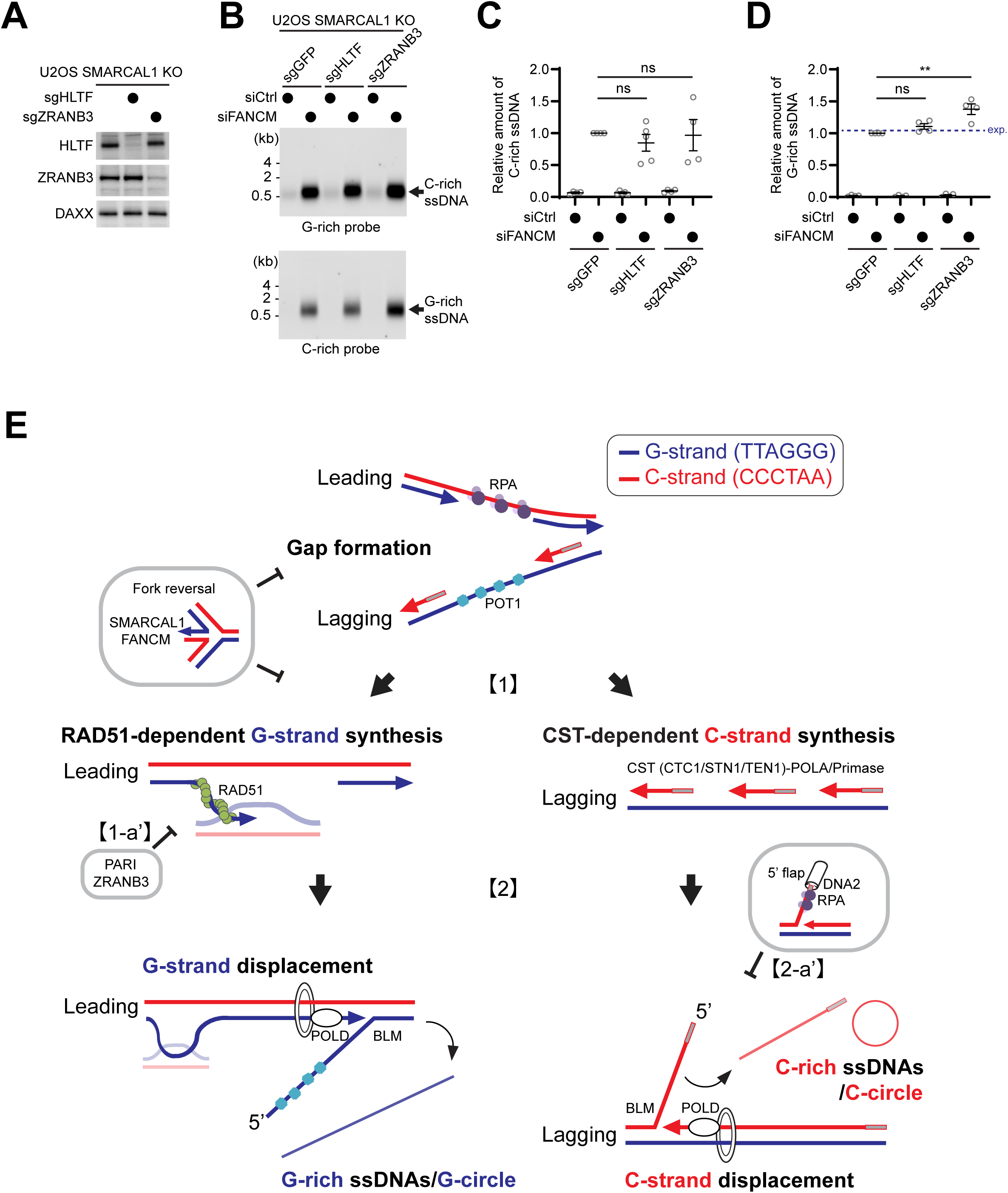
Recombination mediates G-rich ssDNA generation. (A) Western blot analysis of HLTF, ZRANB3, and DAXX (loading control) in U2OS SMARCAL1 KO cells after infection with sgRNAs targeting HLTF, or ZRANB3. (B) 4SET assay with U2OS SMARCAL1 KO cells after infection with sgRNAs targeting HLTF, ZRANB3, and GFP (control sgRNAs). (C) Quantification of the assay in (B); as the relative amount of C-rich ssDNA. (mean ± SEM; unpaired t test). (D) Quantification of the assay in (B); as the relative amount of G-rich ssDNA. Blue dashed line represents the expected value for the siFANCM/sgZRANB3/SMARCAL1 KO condition. (mean ± SEM; unpaired t test). (E) Working model: two distinct mechanisms of C-and G-circle generation in human ALT cancers. Co-deficiency of SMARCAL1 and FANCM induces gap formation at telomeres on both leading and lagging strands. Upon gap formation, G-rich ssDNAs are generated from RAD51-dependent G-strand synthesis on the leading strand, whereas CST-dependent C-strand synthesis is initiated on the lagging strand [1]. PARI and ZRANB3 suppress the formation of G-rich ssDNA [1-a’]. And both strands are engaging in excessive strand displacement of daughter strands which results in C-and G-circle formation [2]. DNA2-mediated 5’ flap processing selectively influences the generation of C-rich ssDNA [2-a’].

Interestingly, ZRANB3 depletion in U2OS SMARCAL1 KO cells significantly increased G-rich ssDNA generation (Fig. 5B, D), an effect not observed upon HLTF depletion (Fig. 5B, D). Neither HLTF nor ZRANB3 depletion altered C-rich ssDNA generation (Fig. 5B-C). In addition to ZRANB3’s role in fork reversal, it resolves recombination intermediates (Ciccia et al. 2012), suggesting the idea that the increased G-rich ssDNA in ZRANB3-depleted SMARCAL1 KO cells might result from its anti-recombination activity.

To further test the role of anti-recombinases, we examined potential human homologs of yeast Srs2: FBH1, RECQ5, and PARI, which we previously reported (Fig. S5D) (Hu et al. 2007; Fugger et al. 2009; Moldovan et al. 2012). We introduced crRNAs targeting FBH1, RECQ5, or PARI genes in U2OS SMARCAL1 KO cells. Depletion of RECQ5. FBH1, and PARI genes were confirmed by Western blot (Fig. S5E), and Quantitative PCR (Fig. S5F), and increased RAD51 filament formation was observed in each gene-depleted condition (Fig. S5G). Consistent with our previous results using PARI siRNA (Fig. 4A-C), PARI depletion significantly increased G-rich ssDNA generation (Fig. S5H, J) while reducing C-rich ssDNA levels (Fig. S5H-I). Despite increased RAD51 filament formation upon FBH1 or RECQ5 depletion, elevated G-rich ssDNA generation was specifically observed only in cells depleted of PARI and ZRANB3, but not FBH1 or RECQ5. Collectively, these findings support the hypothesis that G-rich ssDNA generation is actively suppressed by anti-recombinases such as PARI and ZRANB3, through their inhibitory effects on RAD51-dependent recombination.

## Discussion

G-rich ssDNAs (circular and linear) are present in ALT cancer cells at much lower abundance compared to C-rich ssDNAs; however, the mechanisms underlying G-circle generation have remained unclear, while the mechanism for C-circle generation was recently proposed (Henson et al. 2009; Cesare and Reddel 2010; Jiang et al. 2024; Lee et al. 2024; Lu et al. 2024). Here, we successfully generated abundant G-rich ssDNAs under conditions of SMARCAL1 and FANCM (or FAAP24) co-deficiency in ALT cells. Notably, G-rich ssDNAs (500–3000 nt) are longer than C-rich ssDNAs (200–1500 nt) and predominantly exhibit a more linear structure, distinguishing their unique feature.

Both SMARCAL1 and FANCM function as translocases involved in DNA replication fork reversal. Co-deficiency of SMARCAL1 and FANCM led to hyperphosphorylation of RPA at Serine 33, indicative of potential gap formation. This gap formation resulting from SMARCAL1 and FANCM co-deficiency triggered synergistic increases in both C-rich and G-rich ssDNA generation. Interestingly, generation of both C-rich and G-rich ssDNAs depends on BLM/POLD activity, suggesting that their formation is mediated by strand displacement on daughter strands (See working model in Fig. 5E).

Increased C-rich ssDNA generation under SMARCAL1 and FANCM co-deficient conditions depends on CST complex-mediated C-strand synthesis and is suppressed by DNA2-mediated 5’ flap processing, consistent with canonical C-rich ssDNA generation in ALT cells (Lee et al. 2024). However, G-rich ssDNA generation is independent of both the CST complex and DNA2. We reason that this distinction arises because C-rich ssDNAs originate from lagging strand telomeres, where G-rich sequences are their parental template strand. The CST complex is recruited by POT1, which binds specifically to G-rich telomeric sequences, thereby directing the CST complex to parental lagging strand telomeres (Zhang et al. 2019b; Cai and de Lange 2023). Conversely, G-rich ssDNAs originate from leading strand telomeres, whose parental template strands contain C-rich telomeric sequences, thus no POT1 binding and no recruitment of the CST complex.

In lagging strand telomeres, 5’ flap structures can be efficiently processed by DNA2, as C-rich sequences in these flaps bind RPA, facilitating DNA2 recruitment (Zhou et al. 2015). However, in leading strand telomeres, displaced flap structures contain G-rich sequences that preferentially bind POT1 (Arnoult et al. 2009), competing with RPA binding and potentially hindering DNA2 recruitment. This likely explains the selective inhibition of C-rich ssDNA formation, but not G-rich ssDNA formation, by DNA2.

Here, we identify RAD51 recombinase as essential for initiation of G-rich ssDNA generation. Depletion of RAD51 significantly reduces G-rich ssDNA generation, whereas depletion of PARI, an inhibitor of RAD51 activity, enhances G-rich ssDNA generation. These results suggest that RAD51-dependent gap-filling synthesis initiates G-strand synthesis, leading to the formation of G-rich ssDNAs. Furthermore, ZRANB3, a SNF2-family translocase known to disrupt RAD51-filament intermediates (Ciccia et al. 2012), also suppresses G-rich ssDNA generation.

In contrast, depletion of RAD51 increases C-rich ssDNA generation, likely due to enhanced gap formation, triggering CST complex-mediated C-strand synthesis. This aligns with previous reports showing increased gap formation upon RAD51 depletion or inhibition (Hashimoto et al. 2010; Kolinjivadi et al. 2017; Mann et al. 2022).

Collectively, our findings indicate that both C-circles and G-circles are generated through strand displacement mediated by BLM and POLD at ALT telomeres. However, their specific generation mechanisms, sizes, and structural characteristics differ significantly due to sequence differences and their distinct origins on lagging versus leading strands. Several outstanding questions remain, such as the role of R-loop formation in generating G-rich and C-rich ssDNAs. Notably, RNA/DNA helicases DDX39A and DDX39B have recently emerged from Telomeric ALT *in situ* localization screen (TAILS; a native FISH-based optical CRISPR screening) as potential factors regulating telomeric ssDNA generation, and transcriptional inhibition has also been linked to telomeric ssDNA accumulation (Azeroglu et al. 2025). The effects of transcription and R-loop formation at telomeres on C-circle and G-circle generation are important remaining questions that require further investigation. This is particularly relevant given that FANCM (and its yeast homolog Mph1/Fml1), SMARCAL1, and ZRANB3 repress the formation of R-loops as branchpoint translocases (Silva et al. 2019; Hodson et al. 2022). Additionally, recent CRISPR screening studies have identified synthetic lethality between SMARCAL1 and FANCM due to the accumulation of DNA breaks within repetitive sequences in non-ALT cells (Feng et al. 2024; Fielden et al. 2025). It will be important to determine how repeat instability under SMARCAL1 and FANCM co-deficient conditions resembles or differs from the mechanisms generating C-rich and G-rich ssDNAs at ALT telomeres. Addressing this will further clarify the roles of these translocases in maintaining genomic stability, particularly within repetitive sequences.

## Materials and Methods

### Cell culture

U2OS, and U2OS SMARCAL1 KO cells were maintained in Dulbecco’s Modified Eagle Medium (DMEM) supplemented with 10% fetal bovine serum (FBS), Penicillin-Streptomycin (Gibco, 15070063). G292 cells were maintained in McCoy’s 5a Medium supplemented with 15% FBS. NY cells were maintained in Minimum Essential Medium (MEM) supplemented with 10% FBS. To prevent mycoplasma contaminations, we regularly used Plasmocin Prophylactic (InvivoGen) to maintain the cells according to the manufacturer’s protocol. Cells were subjected to the following treatments with specific concentrations and durations: B02 (27 μM, 48hr), doxycycline treatment (1 μg/ml, 72 hr for G292, NY, U2OS SMARCAL1 KO cells expressing pCW-vector system; pCW-SMARCAL1 WT and R764Q, and pCW-DAXX). Negative control (mock) samples were untreated.

### siRNA-mediated knockdown

siRNA oligos were purchased from Sigma-Aldrich (10 nmol) and purified by desalting. The dTdT overhangs were added to ensure the stability of siRNA. The siRNAs were transfected into the cells at a final concentration of 10 nM, along with Lipofectamine RNAimax (Thermo scientific) according to the manfacturer’s protocol.

The following siRNAs were used:

siBLM: AUCAGCUAGAGGCGAUCAA,

siControl (Universal negative control #1; Sigma-adrich),

siDNA2 (#1+2): CAGUAUCUCCUCUAGCUAGUU; AUAGCCAGUAGUAUUCGAUUU, siFANCM: AAGCUCAUAAAGCUCUCGGAA,

siPARI: AGGACACAUGUAAAGGGAUUGUCUA, siRAD51: GGGAAUUAGUGAAGCCAAA,

siSLX4 (#1+2): GCACCAGGUUCAUAUGUA; GCACAAGGGCCCAGAACAA,

siSTN1: GCUUAACCUCACAACUUAA

### Lentiviral shRNA-mediated knockdown and Cas9/sgRNA-and Cas12a/crRNA-mediated knockout

Lentiviruses were generated by transfecting 293T cells with shRNA vectors targeting human POLD3, STN1, or a control vector; sgRNAs targeting human FANCM, HLTF, or ZRANB3 inserted into pKLV2.2-h7SKgRNA5(SapI)-hU6gRNA5(BbsI)-PGKpuroBFP-W (Addgene #72666); crRNAs targeting human FBH1, RECQ5, or PARI inserted into lenticrRNA hygro vector; pLenti-opCas12a-6xNLS (Addgene #209022); or lentiCas9-Blast (Addgene #52962), along with packaging plasmids pMD2.G and psPAX. Transfections were performed using TransIT (Mirus) according to the manufacturer’s instructions. After 24 hours, the medium was replaced with fresh medium, and cells were incubated for an additional 24 hours. Lentivirus-containing supernatants were collected, filtered through a 0.45 μm syringe filter, and concentrated using a 4x virus concentrator solution comprising phosphate-buffered saline (PBS, pH 7.2), 1.2 M sodium chloride (NaCl), and 40% (w/v) PEG-8000.

The concentrated viral pellets were then used to infect target cells in the presence of 0.5 μg/ml polybrene to enhance viral transduction efficiency. Following infection, stable cell lines expressing Cas9, opCas12a, shRNA, sgRNA, or crRNAs were established through selection with 5 μg/ml Blasticidin-S (Invivogen) for two weeks, 5 μg/ml Puromycin (Invivogen) for 5 days, 200 μg/ml G418 (Invivogen) for two weeks, or 400 μg/ml hygromycin for two weeks. U2OS and U2OS SMARCAL1 KO cells expressing Cas9 were cloned, and clones exhibiting robust Cas9 expression were selected to ensure efficient Cas9/gRNA-mediated gene editing. All experiments involving lentiviruses adhered strictly to proper biosafety protocols.

### Plasmid

**crRNA vector cloning:** To generate the lenticrRNA hygro vector, the U6 promoter-to-crRNA backbone was amplified from plasmid pMV_AA089 (Addgene #216105) using primers F-kpn1-crRNA (taaggtaccgggcagagagggcctatttcc) and R-Nhe1-crRNA (cgccaagcttgctagcgaattc). The resulting PCR amplicon and the lenti-sgRNA hygro (Addgene # 104991) vector were digested with KpnI and NheI enzymes and subsequently ligated. The resulting lenticrRNA hygro vector can be used for crRNA cloning through BsmBI-v2 digestion.

For crRNA cloning, the following sequence (Esmaeili Anvar et al. 2024) was synthesized by IDT:

AGGCACTTGCTCGTACGACGcgtctcgAGATnnnnnnnnnnnnnnnnnnnnnnnTAATTTCTACTATTGTAGATnnnnnnnnnnnnnnnnnnnnnnnAAATTTCTACTCTAGTAGATnnnnnnnnnnnnnnnnnnnnnnnTAATTTCTACTGTCGTAGATnnnnnnnnnnnnnnnnnnnnnnnTTTTTTGAATggagacgTTAAGGTGCCGGGCCCACAT

(’n’ indicates crRNA sequences; four crRNAs per gene were selected from CRISPick, Broad Institute.)

This synthesized crRNA sequence was amplified using primers F-crRNA-gBlock (AGGCACTTGCTCGTACGACG) and R-crRNA-gBlock (ATGTGGGCCCGGCACCTTAA), digested with BsmBI-v2, and ligated into the lenticrRNA hygro vector.

**gRNA vector cloning:** To generate gRNA-expressing vectors, two gRNAs per gene were cloned into the pKLV2.2-h7SKgRNA5(SapI)-hU6gRNA5(BbsI)-PGKpuroBFP-W vector

(Addgene #72666). One gRNA was expressed under the hU6 promoter and the other under the h7SK promoter (Tzelepis et al. 2016).

The oligonucleotides for hU6-gRNA were synthesized as follows:

hU6-gRNA Sense: CACCGnnnnnnnnnnnnnnnnnnn hU6-gRNA Anti-sense: AAACnnnnnnnnnnnnnnnnnnnC

The oligonucleotides for 7SK-gRNA were synthesized as follows:

7SK-gRNA Sense: CTCGnnnnnnnnnnnnnnnnnnn 7SK-gRNA Anti-sense: AACnnnnnnnnnnnnnnnnnnnC

(’n’ indicates the specific gRNA sequences;’n’ in the antisense oligo should be complementary to the sense oligo.)

Sense and anti-sense oligos were phosphorylated using T4 Polynucleotide Kinase (PNK) to add a 5’ phosphate, followed by annealing in a heat block. The annealed hU6-gRNA oligos were ligated into the BbsI-digested pKLV2.2-h7SKgRNA5(SapI)-hU6gRNA5(BbsI)-PGKpuroBFP-W vector. Subsequently, the resulting construct was digested with SapI, and the annealed 7SK-gRNA oligos were ligated into the vector.

### Oligonucleotide and gBlock sequences for gRNA and crRNA generation

U6_S_sgFANCM_1: CACCGATCCACAGGGCGCCCGCGG, U6_AS_sgFANCM_1: AAACCCGCGGGCGCCCTGTGGATC 7SK_S_sgFANCM_2: CTCGAAATTGTACATGACCACGG, 7SK_AS_sgFANCM_2: AACCCGTGGTCATGTACAATTTC, U6_S_sgHLTF_1: CACCGAAAGAAAGTTGGATATGAG, U6_AS_sgHLTF_1: AAACCTCATATCCAACTTTCTTTC, 7SK_S_sgHLTF_2: CTCGTTGGACTACGCTATTACAC, 7SK_AS_sgHLTF_2: AACGTGTAATAGCGTAGTCCAAC, U6_S_sgZRANB3_1: CACCGGGTCTAGGAAAGACAATCC, U6_AS_sgZRANB3_1: AAACGGATTGTCTTTCCTAGACCC, 7SK_S_sgZRANB3_2: CTCGGAAAGACAATCCAGGCAAT, 7SK_AS_sgZRANB3_2: AACATTGCCTGGATTGTCTTTCC, crFBH1: *AGGCACTTGCTCGTACGACG***cgtctcgAGAT**ACCAGACTCCGCAGAGAGCCGAG<u>TAA</u> <u>TTTCTACTATTGTAGAT</u>CCTTCCTCCCGGTGGAAGACCTC<u>AAATTTCTACTCTAGTAG</u> <u>AT</u>TATGCCACAGTTAGACAGGATGC<u>TAATTTCTACTGTCGTAGAT</u>GGTTCTCTCTACA GGTCAGGGAATTTTTT**GAATggagacg***TTAAGGTGCCGGGCCCACAT,* crRECQ5: *AGGCACTTGCTCGTACGACG***cgtctcgAGAT**TGATCCCTATGGGAACCTGAAGG<u>TAAT</u> <u>TTCTACTATTGTAGAT</u>GGGTTAGCAAGTGGTCCACTTGG<u>AAATTTCTACTCTAGTAGA</u> <u>T</u>CCAATGGGGGCATGACTTTCGTC<u>TAATTTCTACTGTCGTAGAT</u>CTGTGCAGAGAGC TTCGAGTTCATTTTTT**GAATggagacg***TTAAGGTGCCGGGCCCACAT,* crPARI: *AGGCACTTGCTCGTACGACG***cgtctcgAGAT**AACATGTCACACAACCATACCAA<u>TAATT</u> <u>TCTACTATTGTAGAT</u>ATCACCTACTGCATACTGTTTGC<u>AAATTTCTACTCTAGTAGAT</u>AA TCTGCTAGTGAATTCAAAGAA<u>TAATTTCTACTGTCGTAGAT</u>TCTCGAGCAGCATGTTT CAAATCTTTTTT**GAATggagacg***TTAAGGTGCCGGGCCCACAT*

### Doxycycline-inducible SMARCAL1 and DAXX expression vector cloning

The doxycycline-inducible expression vector (pCW) was derived from the pCW-Cas9 vector (Addgene #50661). To generate the SMARCAL1 and DAXX inducible expression vectors, the Cas9 coding sequence was removed from the pCW-Cas9 vector by digestion with NheI and BamHI enzymes. SMARCAL1 and DAXX cDNAs were amplified using primers containing NheI and BamHI restriction sites, digested with NheI and BamHI, and subsequently ligated into the digested pCW backbone.

### 4SET (Strand-Specific Southern for Single-Stranded Extrachromosomal Telomeres) DNA Preparation

Cells were harvested using Trypsin-EDTA, pelleted via centrifugation (500 g, 5 min), and washed with PBS. To obtain soluble fractions, cell pellets were resuspended in extraction buffer (PBS containing 5 mM EDTA and 0.1% Triton X-100) and gently pipetted over ten times. The lysate was centrifuged at 1500 g for 15 min to separate the soluble supernatant from the chromatin pellet. To precipitate DNA from the soluble fraction, glycogen (GlycoBlue Coprecipitant, 75 μg) and NaCl (final concentration of 1.6 M) were added, followed by isopropanol (75% of the supernatant volume). DNA precipitation was performed at-20°C for a minimum of 3 hours to maximize recovery of single-stranded DNA fragments. The mixture was centrifuged at 15,000 g for 20 min at 4°C to pellet the DNA. The DNA pellet was washed twice with 70% ethanol, centrifuging at 15,000 g for 5 min each wash. Finally, the DNA pellet was dissolved in 5 mM Tris-HCl (pH 8.0) and incubated overnight at 4°C to ensure complete dissolution. DNA concentration was measured using a Qubit fluorometer (Qubit dsDNA HS Assay Kit, Invitrogen).

### Southern Blot

For electrophoresis, 100 ng of soluble DNA (quantified using Qubit) was loaded into a 0.5% agarose gel and run at 50 V (3.6 V/cm) in TAE buffer (Tris-acetate EDTA). The native agarose gel was transferred onto a positively charged Nylon membrane (Roche) using a vacuum blotting system (VacuGene XL) at 30 mbar for 3 hours in 4×SSC buffer (saline sodium citrate). Subsequently, the membrane was incubated with 1 nM DIG (Digoxigenin)-labeled telomere probes in hybridization solution (DIG Easy Hyb, Roche) at 42°C for overnight. After hybridization, the membrane was washed twice with Wash Buffer 1 (2×SSC, 0.1% SDS) and twice with Wash Buffer 2 (0.5×SSC, 0.1% SDS), each for 10 min. The membrane was blocked in blocking solution (1% Blocking reagent, Roche #11096176001, in maleic acid buffer pH 7.4) for 30 min and then incubated with 0.2% Anti-Digoxigenin-AP Fab fragments (Roche) in blocking solution for 1 hour. Following antibody incubation, the membrane was washed twice in Wash Buffer 3 (maleic acid buffer, 0.3% Tween-20), each for 15 min. Detection was performed by incubating the membrane with 1% CDP-Star substrate (Roche #11759051001) in AP buffer (50 mM Tris pH 9.5, 100 mM NaCl), and signals were detected using the Odyssey Imaging System (LI-COR 2800).

### Western blot

Cell lysates were mixed with sample buffer (375 mM Tris-HCl, pH6.8, 12% SDS, 30% sucrose, 0.012% bromophenol blue, and 30% β-mercaptoethanol) and heated at 55°C for 15 minutes. Samples were then separated on Tris-Acetate or Tris-HCl polyacrylamide gels, and proteins were transferred onto PVDF membranes using the Trans-Blot system. After blocking with either 5% skim milk or 5% BSA in TBS-T (TBS containing 0.1% Tween-20) for 1 hour at room temperature, membranes were incubated overnight at 4°C with primary antibodies. Following washes with TBS-T, membranes were incubated with secondary antibodies for 1 hour at room temperature. After additional washes with TBS-T, membranes were treated with ECL reagent, and protein signals were detected using the Odyssey Imaging System (LI-COR 2800).

### Antibodies

Following antibodies were used for Western blot: anti-actin (Sigma-Aldrich, A5316), anti-ATRX (Santa Cruz, sc-15408), anti-DAXX (Santa Cruz, sc-8043), anti-Lamin A/C (Santa Cruz, sc-376248), anti-STN1 (Abcam, ab89250), anti-SMARCAL1 (Santa Cruz, sc-376377), anti-RAD51 (Santa Cruz, sc-8349), anti-RPA32 (Abcam, ab2175), anti-RPA32 pS33 (Bethyl, A300-246), anti-FANCM (Novus, NBP2-50418; CV5.1), anti-HLTF (Proteintech, 14286-1-AP), anti-ZRANB3 (Thermo, PA565143), anti-RECQ5 (Proteintech, 12468-2-AP), anti-BLM (Bethyl, A300-110), and anti-POLD3 (Proteintech, 21935-1-AP).

### RNA extraction and quantification

Total RNA was isolated using the Direct-zol RNA Miniprep Kit (ZYMO) according to the manufacturer’s instructions. cDNA synthesis was performed using the RevertAid First Strand cDNA Synthesis Kit (Thermo Scientific). Quantitative PCR (qPCR) reactions were carried out with TB Green Premix Ex Taq II (Tli RNase H Plus) (Takara) following the manufacturer’s guidelines.

The following primers were used for qPCR analysis:

GAPDH F: TGAGAACGGGAAGCTTGTCA, GAPDH R: AGCATCGCCCCACTTGATT, SLX4 F: ACTGGGTTTGTGGTGCCAT, SLX4 R: TTGACCATGGCGCCAAAGT, PARI F: AAGGTGCAGCTGCTAGCAA, PARI R: AAGCCACAGCCAGGTCATT, FBH1 F: TGACCCGCTGTTCATTCCT, FBH1 R: ATGCCGTCCACTTTGCTGA

### Immunofluorescence and Image analysis

Cells were fixed in 4% formaldehyde for 10 minutes, permeabilized with PBS containing 0.1% Triton X-100 for 10 minutes, and incubated with primary antibodies in PBS supplemented with 3% (w/v) bovine serum albumin (BSA) and 0.05% Tween 20. The following antibodies were utilized: anti-RPA32 (Abcam, ab2175), anti-RAD51 (Santa Cruz, sc-8349), and anti-TRF2 (Novus, NB110-57130). Samples were imaged at 40X magnification using a Zeiss Axio Imager Z2 microscope. Automated detection of nuclei was performed using MetaCyte software (MetaSystems). RAD51 foci within each nucleus were quantified using the MetaCyte software, whereas RPA32 and TRF2 foci were quantified using ImageJ. Identical exposure times and magnification settings were maintained for all comparative analyses.

### Clonogenic survival assay

To assess clonogenic survival, 300 cells were seeded per well in 24-well plates, followed by a 14-day incubation. Colonies were then fixed and stained with crystal violet solution (0.05% crystal violet, 1% formaldehyde, 1x PBS, 1% methanol).

### Chromatin isolation assay

Cells were lysed in a hypotonic buffer (1% NP-40, 10 mM NaCl, 5 mM MgCl_2_, and 20 mM Tris-HCl, pH 8.0) supplemented with protease and phosphatase inhibitors, then incubated on ice for 1 hour. The lysate was centrifuged at 4 °C, and the supernatant was retained as the cytoplasmic fraction. The pellet was resuspended in a hypertonic buffer (1% NP-40, 150 mM NaCl, 5 mM MgCl_2_, and 20 mM Tris-HCl, pH 8.0) and subjected to sonication. Following centrifugation, the supernatant was isolated as the nuclear fraction.

### Statistical analyses

Statistical analyses for the experiments were conducted using GraphPad Prism 10. The graph displays all individual values. Detailed information regarding the tests and corresponding p-values is provided in the figures and their respective legends.

## Author contribution

A.C., and J.M. conceived the project; Junyeop Lee, A.C., and J.M. designed research; Junyeop Lee, E.J.S., Jina Lee, A.T., and J.M. performed research; Junyeop Lee, E.J.S., Jina Lee, A.T., A.C., and J.M. analyzed data; and J.M. wrote the paper with input from all authors.

## Acknowledgement

We thank Dr. Alan D’Andrea for suggesting the FANCM experiment, Dr. Vincenzo Costanzo for helpful discussions on SMARCAL1 function, Dr. Ralph Scully for insightful discussions regarding DNA recombination, and Dr. Greg Ira for valuable discussions on DNA2 function. We also greatly appreciate Drs. Jean Gautier and Max Gottesman for their valuable discussions, advice, and support. This work was supported by NIH grants R35GM155138 (J.M.) and R01CA197774 (A.C.).

